# Chemoproteomic profiling of serine hydrolases reveals the dynamic role of lipases in *Phaeodactylum tricornutum*

**DOI:** 10.1101/2024.03.12.582592

**Authors:** Achintya Kumar Dolui, Beery Yaakov, Weronika Jasinska, Simon Barak, Yariv Brotman, Inna Khozin-Goldberg

## Abstract

*Phaeodactylum tricornutum* is a model oleaginous pennate diatom, widely investigated for the accumulation of triacylglycerols (TAG) in lipid droplets during nitrogen (N) starvation. However, lipid droplet breakdown, TAG catabolism, and remobilization upon N replenishment during growth restoration are less studied. Serine hydrolases (SH) constitute a diverse family encompassing proteases, amidases, esterases, and lipases. In this report, we adopted a chemoproteomic approach called Activity-Based Protein Profiling (ABPP) to explore the repertoire of active serine hydrolases to elucidate the mechanisms of lipid metabolism in *P. tricornutum* (strain Pt4). A superfamily-wide profile of serine hydrolases revealed a differentially active proteome (activome) during N starvation and after nutrient replenishment. We report 30 active serine hydrolases, which were broadly categorized into metabolic serine hydrolases and serine proteases. Lipases appeared to be the major metabolic linchpins prevalent during lipid remobilization. Global transcriptomics analysis provided a complementary insight into the gene expression level of the detected serine hydrolases. It revealed putative phospholipases as central players in membrane lipid turnover and remodeling involved in cellular lipid homeostasis and TAG accumulation. TAG remobilization and lipid droplet breakdown were impaired in the presence of phenyl mercuric acetate (PMA), whose activity as an SH inhibitor was validated by competitive ABPP. Lipid species profiling corroborated the impairment in TAG degradation and the buildup of structural lipids in the presence of PMA after nutrient replenishment. Collectively, our functional proteome approach, coupled with the transcriptome and lipidome data, provides a comprehensive landscape of *bona fide* active serine hydrolases, including lipases in this model diatom.

## Introduction

Diatoms are an important group of photosynthetic eukaryotes belonging to the Stramenopiles, which have gained global ecological significance as primary producers (Sarthou et al., 2005), and have become important in biotechnology. *Phaeodactylum tricornutum* is a model oleaginous pennate diatom; it has a wide range of biotechnological applications as it produces omega-3 LC-PUFA, high-value carotenoids, and accumulates storage lipids as carbon and energy-rich reserves (Butler et al., 2020). In their natural environment, diatoms rely on responses to environmental cues by diverting carbon and nitrogen (N) into macromolecules to maintain energy balance and cell homeostasis (Wagner et al., 2017). Nitrogen (N) deficiency and some other nutrient imbalances induce triacylglycerol (TAG) accumulation in *P. tricornutum*. TAG sequestration in lipid droplets (LDs) is favored by modulating the cellular metabolism, altering the flux of carbon towards storage lipid synthesis and membrane lipid remodeling (Levitan et al., 2015; Kim et al., 2017). As a result, the photosynthetic apparatus is cannibalized, and growth is severely arrested (Levitan et al., 2015). During recovery from N stress after nutrient resupply, LD breakup and TAG remobilization play a vital role in providing energy and acyl groups for growth resumption and the ensuing resynthesis of structural membrane lipids (Jallet et al., 2020). Storage lipid remobilization ultimately fuels membrane reorganization and restoration of a functional photosynthetic apparatus (Solovchenko, 2012; Tsai et al., 2015). Therefore, LD biogenesis and lipolysis are two distinctive and opposing phenomena that could be investigated by exploring the active proteome involved in lipid metabolism.

Lipases and esterases, the constituent members of the serine hydrolase (SH) superfamily, are the linchpins of lipid catabolism (Faucher et al., 2020). The inhibition of TAG turnover was proposed as a tool to enhance the potential of *P. tricornutum* and other oleaginous microalgae as a cell factory for sustainable TAG production without compromising growth. Since lipid catabolism is not a central metabolic pathway, its targeted inhibition by selectively knocking down lipase-encoding genes would be a rational approach to enhance lipid yield while maintaining optimal microalgal growth (Trentacoste et al., 2013). Previous studies have reported the gene expression profiling of lipases in *P. tricornutum* under nutrient stresses (Alipanah et al., 2015; Remmers et al., 2018). However, relatively little is known regarding the repertoire and function of lipases involved in TAG degradation following N resupply to N-starved cells. A recent study used a bioinformatics approach to decipher a repertoire of lipolytic enzymes in *P. tricornutum* (Murison et al., 2023). Two TAG lipases have been functionally characterized in *P. tricornutum*, namely TGL1, which shares protein sequence similarity with *Arabidopsis thaliana* SDP1 (Barka et al., 2016), and OmTGL, which is predicted to localize to the outermost membrane of the complex plastid (Li et al., 2018). However, the functional annotation and validation of the putative lipid hydrolases in *P. tricornutum* is challenging, since diatom proteins may not share an apparent homology with well-known proteins described in plants, animals, or other organisms (Leyland et al., 2020). Indeed, identification of the active SHs, especially the lipases involved in lipid metabolism, is largely lacking in microalgae. Enumerating the cryptic and active enzymes will have a significant impact on the fundamental understanding of the mechanisms of TAG metabolism and LD turnover in microalgae, and for genetic engineering of this diatom.

In this study, we adopted a functional proteome approach called Activity-Based Protein Profiling (ABPP) - the brainchild of Cravatt and colleagues (Liu et al., 1999) - for investigating active SHs in *P. tricornutum.* The hallmark of ABPP is its functional readout of reactive enzymes rather than being limited to protein or mRNA abundance (Speers and Cravatt, 2009). This powerful functional proteomics technology, coupled with advanced mass spectrometry, has revolutionized the concept of proteomics research. It is an excellent method of choice for investigating the active proteome involved in the dynamic LD metabolism of *P. tricornutum*. Moreover, we complemented the functional SH proteome with its primary biochemical output (lipidome) during recovery from N starvation. We also performed global transcriptome profiling to monitor changes in the mRNA abundance of genes encoding enzymes/proteins involved in lipid metabolism under, and following recovery, from N starvation.

## Results

### Labeling of SHs and competitive gel-based ABPP

First, *in vitro* gel-based ABPP was performed to obtain an initial assessment of functional SHs in N-starvation and after N resupply. A fluorophosphonate (FP) serine hydrolase probe was used to label protein samples processed from N-depleted cells (-N (120 h)) designated as the starvation sample, and from N-depleted cells that were replenished with nutrients (+N (24 h)) designated as the recovery sample. To achieve optimal labeling of serine hydrolases, a Tris buffer (pH 8.0) was chosen for extracting proteins in the native state with proper folding (Speers and Cravatt, 2009; Zweerink et al., 2017). The probe-labeled profile of the proteome on SDS-PAGE under N-starvation conditions differed from that of the recovery samples **(Figure 1A, 1B)**. The signal around 40 kDa was more prominent in N-starvation than that of the recovery sample. The signal of another differentially active protein showed an opposite trend whose intensity got stronger during the course of recovery (**Supplemental Figure S1A**). However, a predominant signal around ∼90 kDa was present both during N-starvation and recovery conditions, albeit with a stronger intensity in the proteome of the recovery samples (**Figure 1A, B**). The in-gel fluorescence profile provided us a snapshot of the activity of SHs under N starvation and 24 h after N resupply.

**Figure 1.**
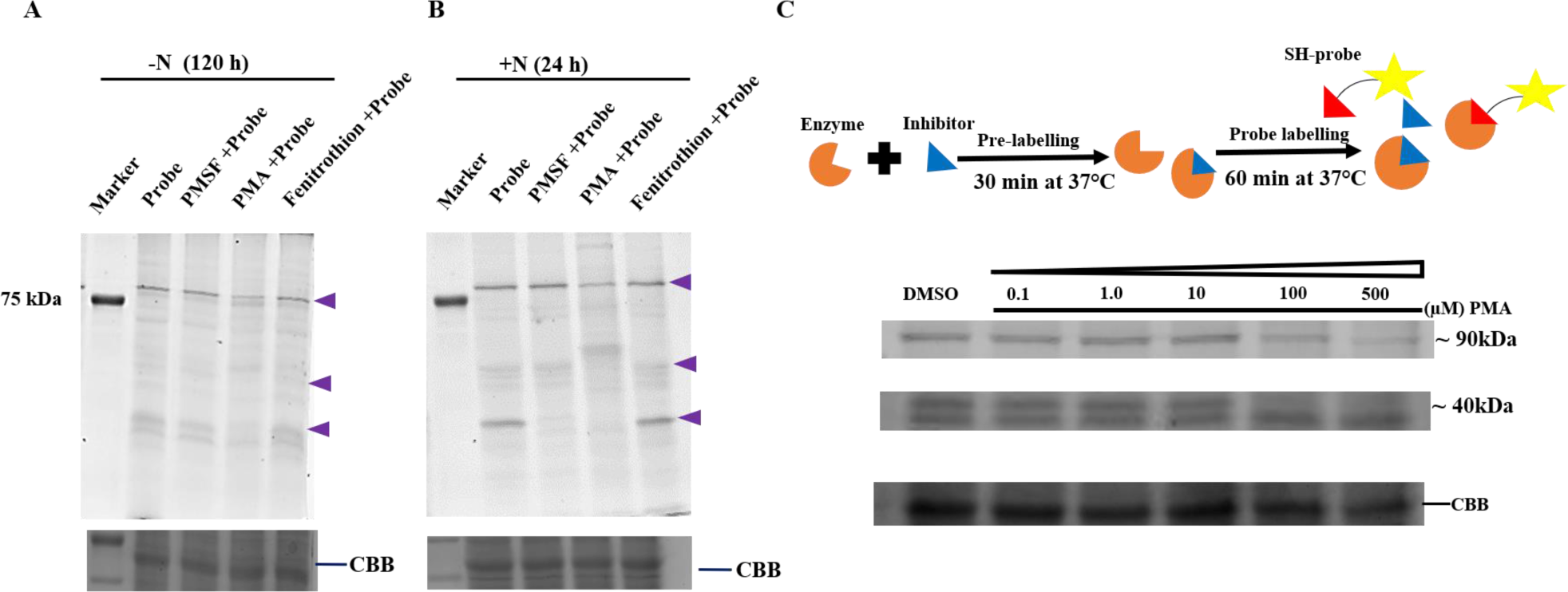
Gel-based competitive ABPP using a FP-probe displays a differential labeling pattern in N-starvation and recovery samples. For this assay, 30 µg protein, 100 µM of indicated inhibitors, and 2 µM probe were used. Samples were preincubated with inhibitors at 37 °C for 30 min followed by labeling with the probe for 60 min at 37 °C; (A) Fluorescent readout of proteome processed from the N-starved cells (120 h) and the cells 24 h after N replenishment (B), respectively. A protein marker depicts a band of 75 kDa detectable due to its fluorescent signal. The arrow points to ablated signal in PMA and/or PMSF pre-treated samples and indicates a potential serine hydrolase. (C) A schematic representation of the competitive ABPP assay and a concentration dependent inhibition of 24 h-recovery proteome with phenyl mercuric acetate (PMA). A representative image of Coomassie Brilliant Blue (CBB) stained gels for each set of gel-based ABPP experiments is included as a loading control.

Gel-based competitive ABPP can be used to screen uncharacterized enzymes in their native forms with specific inhibitors, as no *a priori* knowledge of the substrate is required (Simon and Cravatt, 2010). Three inhibitors were used: phenylmethylsulfonyl fluoride (PMSF), fenitrothion, and phenyl mercuric acetate (PMA), against the probe to screen the proteome of the N-starvation and 24-h recovery samples. PMSF has been shown to be a potent and covalent SH inhibitor in competitive ABPP in bacterial (Ortega et al., 2016) and plant systems (Dolui and Vijayaraj, 2020). Fenitrothion has been shown to inhibit fatty acid amide hydrolase (FAAH), as well as monoacylglycerol lipase (Suzuki et al., 2013), while PMA was reported as a lipase and esterase inhibitor (O’Sullivan et al., 1987; Liederer and Borchardt, 2006). A substantial reduction (∼50%) in the signal intensity around ∼90 kDa was observed in the sample pre-incubated with PMA (**Fig. 1 A, B**). There was also a substantial attenuation of the signal around ∼40 kDa in the sample pre-incubated with PMSF (**Fig. 1B**). Importantly, this signal intensity was also compromised by PMA, indicating that it derives from a *bona fide* SH. Since strong signal reduction was observed in the proteome pre-incubated with PMA, a concentration-dependent competitive ABPP was further performed using a range of PMA concentrations. While the signal ∼40 kDa was completely abolished at higher PMA concentrations (100 µM and 500 µM), the signal ∼90 kDa was ablated by half at higher PMA concentrations **(Fig. 1C)**.

### In vivo inhibition of lipolytic activities by PMA

Next, we tested a range of PMA concentrations in *P. tricornutum* cultures during early stages of recovery from N starvation when maximum lipase activity is required for LD breakdown and remobilization of TAG. Three concentrations (25 nM, 50 nM, and 100 nM) were used since the higher PMA concentrations were lethal (**Supplemental Fig. S2**). A dose-dependent growth repression was evident with 50 and 100 nM PMA applied during recovery (**Fig. 2A, B**) and was associated with a delayed breakdown of LDs and recovery of plastids as observed by microscopy (**Fig. 2C**). It should be noted that PMA also inhibited growth in replete (+N) conditions **(Supplemental Fig. S3),** suggesting that it may inhibit lipases involved in TAG homeostasis. To validate impaired TAG degradation, total lipids were extracted and resolved by TLC for semi-quantitative analysis of TAG. Visualization of TAG spots on TLC plates confirmed a delayed degradation of TAG in cells treated with PMA (**Supplemental Fig. S4**).

**Figure 2.**
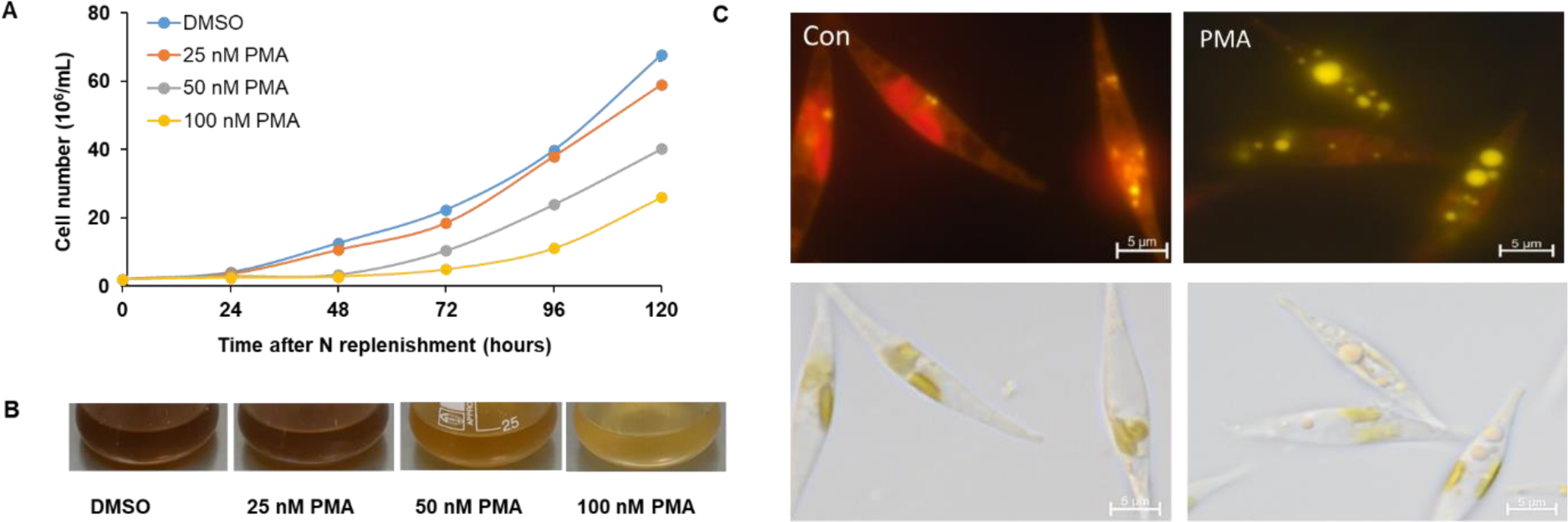
**PMA inhibits lipid droplet degradation and hinders growth resumption in *P. tricornutum* following N replenishment**. Nitrogen-starved cells (2 ·10^6^ mL^-1^) were resuspended in (+N) replete medium and treated with 25, 50 and 100 nM PMA. Cell numbers were monitored over 120 h. Cultures supplemented with DMSO (PMA solvent) served as a control. (**A**) Growth of control and PMA-treated cultures; (**B**) *P. tricornutum* cultures 72 h after N replenishment; (**C**) Micrographs of *P. tricornutum* cells at 48 h of recovery with and without (DMSO control) 50 nM PMA. Cells were stained with the lipophilic dye Nile Red to visualize bright yellow fluorescence of lipid droplets and red fluorescence associated with membrane lipids.

### Lipidomic analysis to validate the effect of PMA on P. tricornutum glycerolipids after 24h of nutrient resupply to N-deprived cells

Untargeted lipid profiling by LC-MS was performed to assess in more detail the impact of PMA on the *P. tricornutum* lipidome after 24 h of recovery. The principal component analysis (PCA) of all identified lipid species and a volcano plot of the 40 most significantly different species revealed a clear separation of the control and the PMA-treated lipidomes (**Supplemental Fig. 5)**. Most of the TAG species were significantly more abundant in the lipidome of PMA-treated cells compared with untreated cells, including major species with a total of 46, 48, 50, 52 and 56 carbons in acyl chains (**Fig. 3**), reflecting the inhibition of TAG hydrolysis. A significantly higher abundance of 34:2 (18/16), 36:5, 36:6 (20:5/16:0-1) and some other diacylglycerol (DAG) species, as well as of phosphatidylcholine (PC) species 32:1, 32:2 (16/16), 34:5 (20:5/14:0) and a series of 36:4-6 (20:5/16:1) species was also observed in the PMA-treated cells (**Fig. 3**). In contrast, the molecular species of the main plastidial galactolipid monogalactosyldiacylglycerol (MGDG), in particular comprising C16 acyl groups (32:2 – 32:7), were present at a lower abundance in PMA-treated cells compared to control, while the dominant EPA-containing MGDG species 36:8 (20:5/16:3) did not exhibit significant differences. A few species of digalactosyldiacylglycerol (DGDG) 32:1 (16:1/16:0) and 36:6 (20:5/16:1) were present in higher abundances in the PMA-treated cells, while DGDG 36:3 (putatively 18:2/18:1), 38:5 (20:5/18:0), and 38:6 (20:5/18:1) species were significantly higher in the control. Sulfoquinovosyldiacylglycerol (SQDG) species of 32C series (16/16) and 34 (18/16) were significantly elevated in the untreated control as well. The overall pattern of changes in TAG and polar lipid species confirmed that PMA hindered storage lipid remobilization and remodeling and restoration of membrane lipids during recovery, likely due to the inhibition of lipid hydrolases, in agreement with the results of gel-based ABPP.

**Figure 3.**
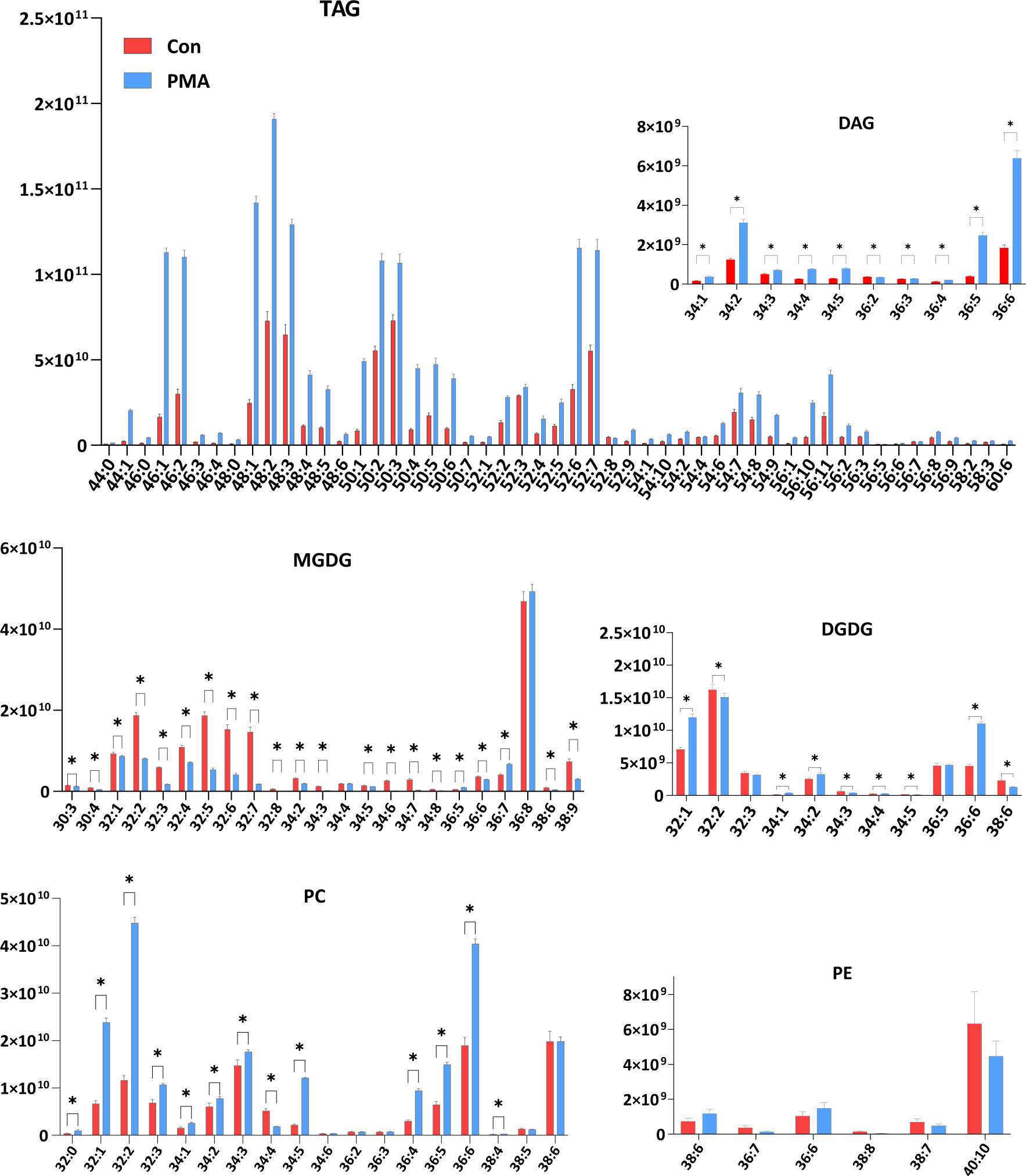
Effect of PMA on the glycerolipidome of *P. tricornutum* after 24 h of recovery from N starvation. Relative normalized intensities of TAG and polar lipid species are indicated. Data are mean ± S.D. (n = 4). Asterisks indicate significant difference at p < 0.05 (students t-test). Note: all species of TAG except 52:8 showed significant differences and were therefore not labeled.

### Enrichment and identification of active SHs by gel-free ABPP

For global assessment of SH activities, gel-free ABPP was performed with an FP- desthiobiotin probe using on-bead digestion and active site peptide profiling according to Dolui and Vijayaraj (2020). Thirty SHs were enriched and reproducibly present in samples of multiple biological replicates of on-bead digestion and/or sustained through stringent selection criteria in active site peptide profiling (**Fig. 4A and B**). The identified SHs were classified into metabolic serine hydrolases (lipases/esterases, amidases, protein and glycan hydrolases) and serine proteases (trypsin/chymotrypsin/subtilisin enzymes). SHs were further categorized into different catalytic members of this superfamily based on their existing annotation and functional domains and motifs **(Fig. 4C**). Approximately 54% of the enriched SHs were found to be metabolic serine hydrolases (mSHs). Eleven mSH activities were detected during recovery from N-starvation and eight of them are predicted to have lipolytic activity (**Fig. 4D**). Enrichment of certain SHs exclusively in active-site peptide profiling was a striking feature of the gel-free ABPP approach. These low copy number proteins that could evade detection in the on-bead digested sample were detected with active site peptide profiling (**Supplemental File S1**). By integrating these two platforms of gel-free ABPP, a consolidated list of active SHs was generated regardless of their relative abundance in the proteome (**Fig. 4B**).

**Figure 4.**
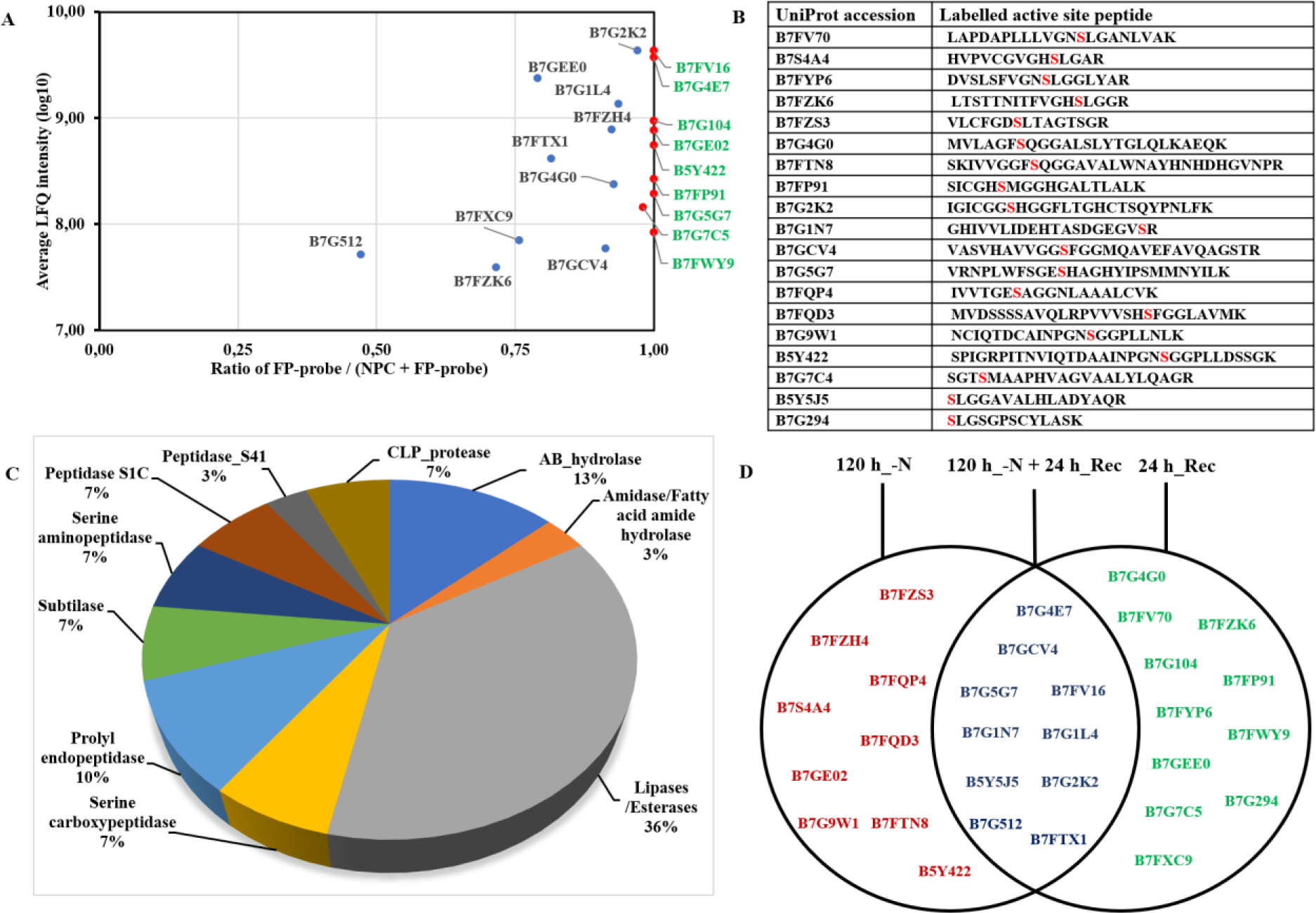
Enrichment of the functional SHs by gel-free ABPP using FP-desthiobiotin probe. *P. tricornutum* cell-free extracts were labeled with and without probe (DMSO), and proteins were enriched on streptavidin agarose resin beads followed by trypsin digestion (*On-bead digestion*). Alternatively, the labeled proteins were digested in solution followed by enrichment of digested peptides (*Active site peptide profiling*) on streptavidin agarose. The digested peptides were subjected to LC-MS analysis. (**A**) Distribution of the probe labeled (FP) on-bead digested proteins over the no-probe control (NPC: DMSO). The average LFQ scores were plotted against the distribution of the LFQ scores for each protein detected with and without the probe. All red spots designate highly enriched proteins and are highlighted on the right. (**B**) The table represents the active SHs identified by active site peptide profiling. UniProt accession IDs for each protein are followed by their corresponding active site peptides. The active site serine modified by the labeling of the probe, is highlighted in red. (**C**) Distribution of the identified SHs into different catalytic members of this superfamily. (**D**) A pictorial representation of the differential activome for the identified SHs. The proteins represented by red and green colors are found exclusively in the N-starvation and recovery samples, respectively. Blue color highlights proteins shared in both N starvation and recovery samples.

### Gel-free ABPP uncovers lipases prevalent during recovery

One of the overarching goals of this study was to uncover the active lipid hydrolases involved in TAG homeostasis and TAG degradation during recovery from N starvation. We have successfully identified 11 lipases/esterases, which were enriched using fluorophosphonate probe, FP-desthiobiotin. These represented ∼70% of the total mSHs identified in this study and can possibly hydrolyze a wide range of lipid substrates. None of these lipases were previously characterized, underscoring the potential of ABPP as a tool of choice to unravel these lipid catabolic linchpins. A few lipases were found exclusively in recovery samples, for example, B7G4G0 annotated as a putative lysophospholipase. An abundant putative class 3 lipase (B7FV70), which features the conserved active site consensus sequence GxSxG. In addition, as predicted by PHYRE 2, it bears a considerable secondary structure similarity to lysosomal acid lipase (**Fig. 5A**). Another lipase (B7FZK6), a mitochondrial resident protein as predicted by HECTAR (Gschloessl et al., 2008) and WolfPSort, shares a canonical pentapeptide GHSLG sequence present in hormone sensitive as well as fungal lipases (**Fig. 5A**). It has the PGAPI motif found in TGL2 (Tgl2p), the functionally characterized mitochondria-localized TAG lipase in *Saccharomyces cerevisiae* (Ham et al., 2006). A putative carbohydrate esterase (B7G104), which belongs to the SGNH hydrolase superfamily and possesses a Sialate O-acetyl esterase (SASA) domain, was enriched in recovery samples. It may potentially mediate the deacylation of glycolipids in *P. tricornutum*, was also enriched in recovery samples. The identified lipase portfolio also featured two low molecular weight lipases, which were found to be functional exclusively under N starvation: a GDSL lipase (B7FZS3/29.36kDa) of the SGNH hydrolase superfamily and a relatively small phospholipase (B7FTN8/∼27kDa). The identified GDSL lipase possesses a Lipase_GDSL_2 domain in the N terminal domain (**Fig. 5A**). To the best of our knowledge, a functional GDSL lipase from a diatom has not been previously reported.

**Figure 5.**
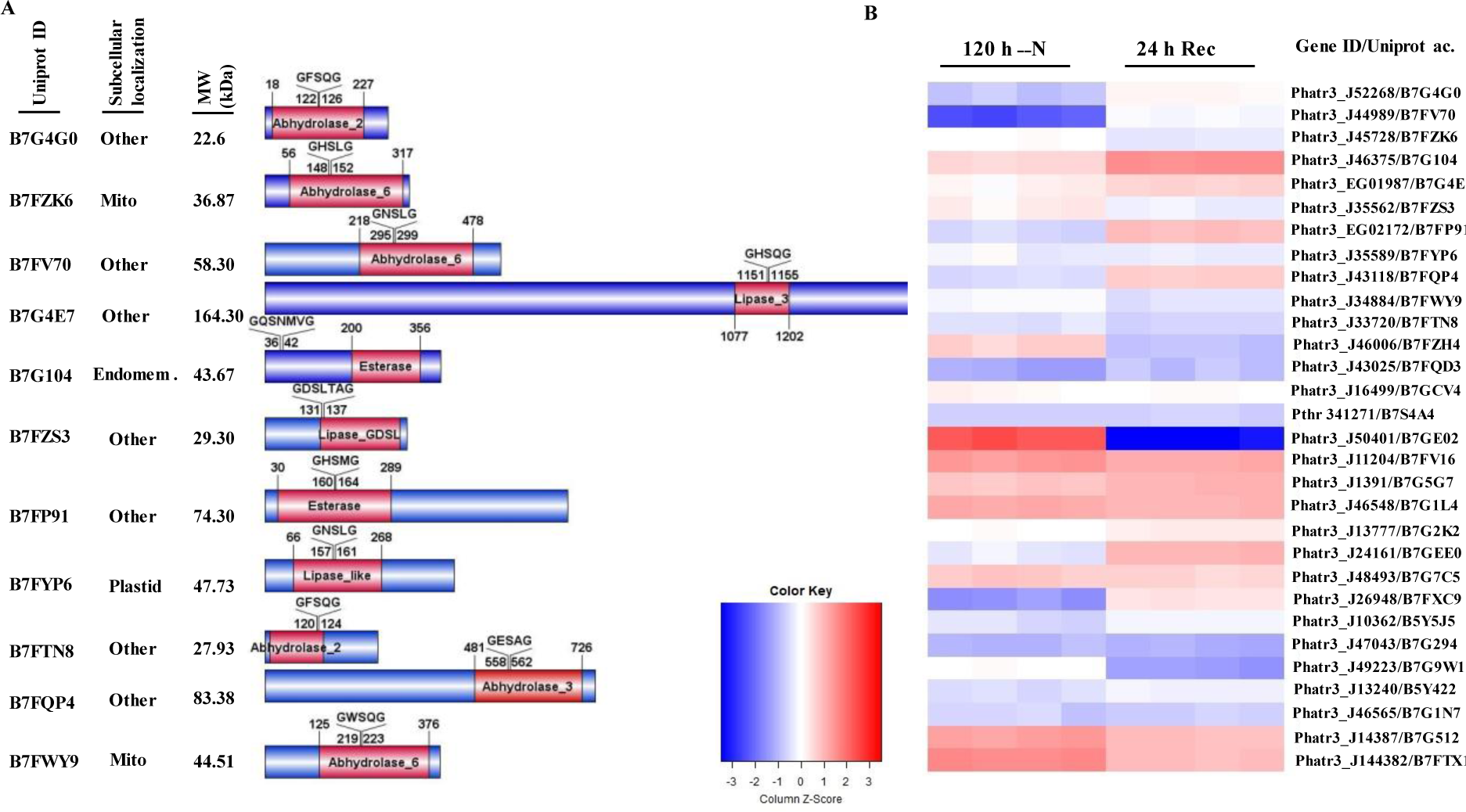
Gel-free ABPP reveals a unique enrichment of functional SHs overrepresented by lipases/esterases. (**A**) List of the detected lipases. Protein accession IDs (UniProt) predicted subcellular localization and predicted molecular weight (MW; kDa) are indicated. The domain organization and the conserved sequence GxSxG in the protein are highlighted. (**B**) Heat-map of relative gene expression values obtained from RNA-Seq of all detected SHs in N-starvation and recovery. Log2-transformed expression values for each gene encoding the identified proteins were used to generate the heat map.

### Enrichment of non-lipid-metabolizing mSHs during N starvation

In addition to lipases/esterases, we report five other metabolic serine hydrolases, constituting approximately 30% of the total mSHs. Two chlorophyllases, one amide and two alpha beta hydrolase domain containing proteins (ABHD). One of the hallmarks of N-starvation is the substantial ablation of plastids. Consistent with this phenomenon, two chlorophyllases (B7S4A4, B7FQD3) were identified, with the former possessing a lipase consensus sequence featuring the catalytic triad of typical lipases. A potential amidase (B7GE02) was found in N-starved samples **(Fig. 5)**, representing one of the predominant SH detected in N-starved cells.

An orphan ABHD-containing SH (B7FZH4) was identified whose role and physiological substrate remain elusive. Another notable finding from the gel-free ABPP was the enrichment of another AB hydrolase superfamily protein (B7GCV4), belonging to the MetX_acytransf family and predicted to have transferase activity along with its prototypical hydrolase activity.

### Enrichment of serine proteases encompassing members of diverse proteolytic pathway

In addition to mSHs, we also report 14 serine proteases (∼46% of identified SHs). In contrast to mSHs, these serine proteases are well annotated, albeit not functionally characterized. Accordingly, we classified them into seven different subclasses, including serine carboxypeptidase/peptidase_S10, prolyl endopeptidase/peptidase_S9, peptidase_S8/Subtilase, serine aminopeptidase/S33, peptidase S1C, Peptidase_S41, and CLP_protease/ClpP_Ser_AS **(Fig. 4C)**. These serine proteases were enriched in both N-starved and recovery samples. However, a few serine proteases were enriched exclusively either in N-starvation or in recovery samples. For example, peptidase S1C (B7G9W1) was found to be functional in N-starvation.

### RNA-seq analysis corroborates the activity of enriched SHs

To gain insight into the interrelation between mRNA abundance and protein activity of enriched SHs, we performed RNA-seq analysis on N-starved and recovery cells. Differential expression analysis showed that out of 11,272 genes, 4,593 genes were upregulated and 4,562 genes were downregulated after 24h of recovery compared to N-starvation (**Supplemental File 2)**. Significantly differentially expressed genes were subjected to Gene Ontology (GO) enrichment analysis, which indicated a substantial revitalization of many essential cellular processes and functions within 24 h after the N resupply (**Supplemental File 3**). Hereafter, we describe the expression patterns of genes encoding SHs revealed in this study.

A good agreement between activity and mRNA level was generally found for most of the SHs identified by ABPP. For example, the significantly upregulated expression of the Phatr3_J46375 gene (adjusted p value = 9.13E-10) encoding esterase (B7G104) during recovery reinforces the assertion that this esterase is functional during this process (**Fig. 5B**). An increased expression of the gene Phatr3_J52268 encoding phospholipase (adjusted p value = 9.67E-15), and of the gene Phatr3_J24161 encoding protease (B7GEE0) was determined exclusively during recovery. Further, a strong mRNA-protein correlation was found for the amidase (B7GE02); the gene encoding this SH (Phatr3_J50401) was significantly upregulated (adjusted p value = 0) during N starvation and downregulated during recovery **(Fig. 5B)**. The enrichment of this amidase by ABPP during recovery could be attributed to its sustained activity, despite the downregulation in its gene expression.

### Genome mining for cataloging putative lipases of *Phaeodactylum tricornutum*

In addition to the chemoproteomic approach, we enlisted 45 lipases using *P. tricornutum* genome portals and resources (Ensemble protists https://protists.ensembl.org, DiatOmicBase https://www.diatomicsbase.bio.ens.psl.eu; UniProt https://www.uniprot.org/) based on their existing annotation. The putative lipases were further classified into different subclasses (class 3 TAG lipases, patatin-like lipases, phospholipases) based on their domain organization and putative substrate specificity (**Fig. 6**). It is noteworthy that the patatin-like protein family also includes characterized plant and microalgal TAG lipases bearing an N-terminal domain of unknown function (DUF3336) adjacent to the patatin domain.

**Figure 6.**
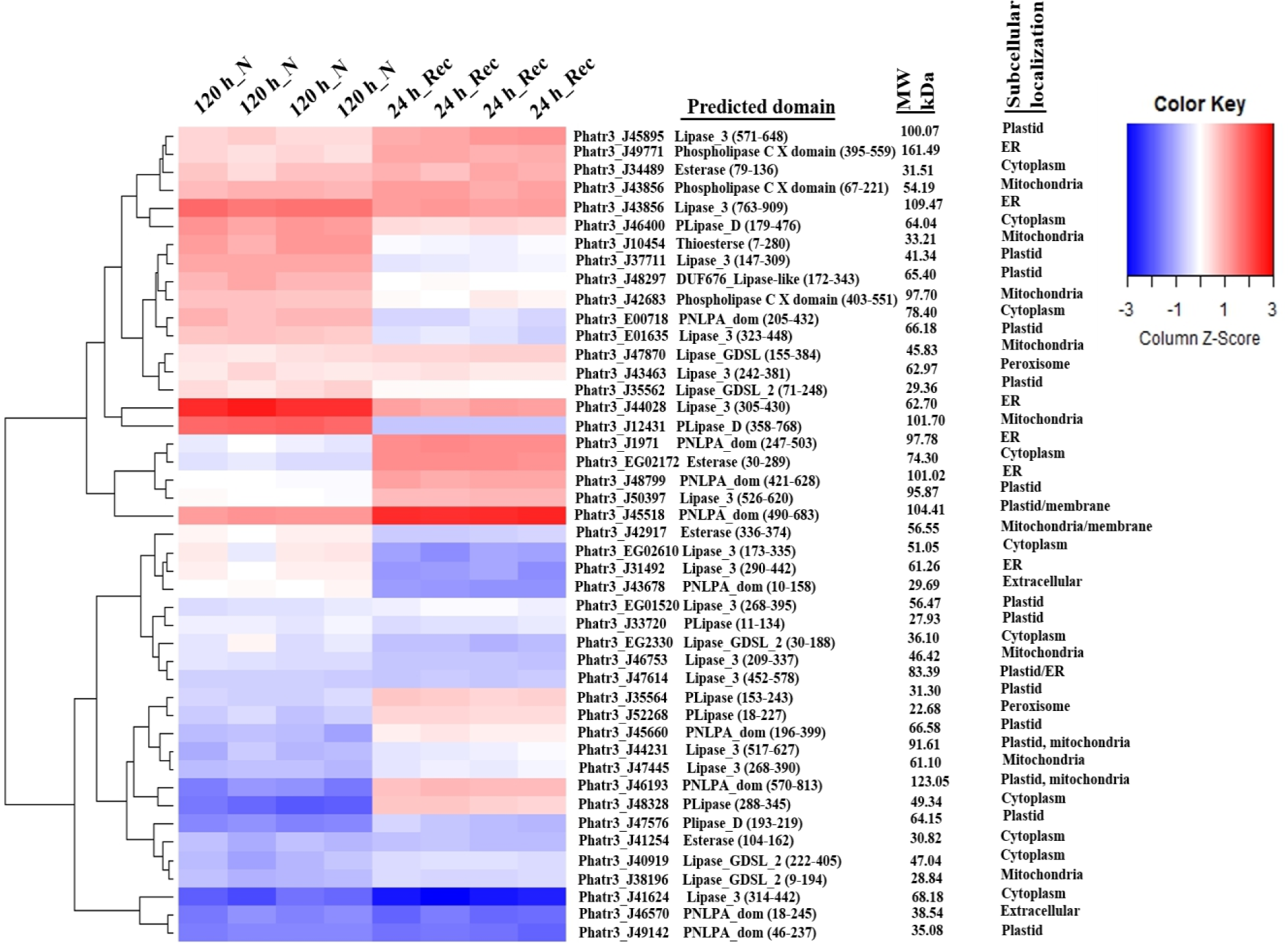
**Expression response of 45 putative lipase genes of *P. tricornutum* to N-starvation and recovery**. The heat map is based on log2-transformed expression values of each lipase gene. Identified putative lipases were classified into GDSL lipases, phospholipases C and D, patatin-like phospholipases, esterases and class 3 lipases based on their existing annotation. Normalized expression values of four replicates of N starvation and four replicates of recovery samples are represented in the heat map. Red color represents higher transcript abundance and blue color represents decreased transcript abundance. The gene ID and the predicted domain of each lipase transcript are given on the right. The length of the predicted domains is indicated in the parenthesis. PNPLA_dom, patatin like phospholipase domain; DUF, domain of unknown function; PLipase_D, Phospholipase D; Lipase_3, Class 3 lipase.

*Phaeodactylum tricornutum* has two TAG lipases of this domain structure, Tgl1 (Barka et al., 2016) and TAG-like B encoded by Phatr3_J1971 and Phatr3_J48799, respectively. Notably, our transcriptome analysis revealed that expression of four genes (Phatr3_J1971(adjusted p value =3.18E-42), Phatr3_J45518 (adjusted p value =1-15E-58), Phatr3_J45660 (adjusted p value =4.20E-32) and Phatr3_J46193 adjusted p value =2.06E-120) encoding patatin-like phospholipases (PNPLA) was upregulated exclusively during recovery (**Fig. 6**). Unlike phospholipase A1 (PLA1) and phospholipase A2 (PLA2), which cleave FAs from the *sn*-1 or *sn*-2 position of the glycerol backbone, respectively, PNPLAs can cleave FAs from both positions of different glycerolipids, but with different specificity (Li et al., 2013).

Phospholipases C and D are phosphodiesterases, which do not belong to the SH superfamily, therefore, these lipases were not enriched in our samples, however, significant insights regarding their roles could be obtained from the transcriptome data. Phospholipase C (PLC) and phospholipase D (PLD) generate the PA and DAG backbone from phospholipids, respectively. Consistent with their possible role in polar lipid mobilization towards TAG biosynthesis during N starvation, expression of the genes encoding phospholipase D (Phatr3_J12431) (adjusted p value = 0) and phospholipase C (Phatr3_J42683) (adjusted p value = 6.85E-38) was significantly upregulated under N starvation **(Fig. 6)**.

### Transcriptome analysis corroborates membrane lipid synthesis, remodeling, and trafficking during recovery from N starvation

To the best of our knowledge, there are no previous reports of transcriptional profiling of lipid metabolism genes during recovery from N starvation in *P. tricornutum*. In the following section, we describe the significant changes in the expression of genes related to lipid metabolism, other than lipases presented in Fig. 6.

In line with the reestablishment of fatty acid and lipid synthesis in the plastid, expression of genes encoding an acyl carrier protein (Phatr3_J9709) and four plastidial triose phosphate translocators (Phatr3_J24610, Phatr3_J50742, Phatr3_J54017, Phatr3_J43320) was upregulated during recovery (**Supplemental File 2**). The expression of genes encoding enzymes mediating the biosynthesis of the plastidial glycerolipids was also upregulated. These genes include MGDG synthase (*MGD1*/Phatr3_J54168), DGDG synthases (Phatr3_J11390, Phatr3_J43116), UDP-sulfoquinovose synthase (*SQD1/*Phatr3_J21201), as well as two plastidial desaturases (FAD6/Phatr3_J48423, PlastidΔ6FAD/Phatr3_EG02619) (**Table 1**). Expression of the gene encoding acyltransferase (LPCAT/Phatr3_J20460), involved in acyl editing and membrane lipid remodeling (Jasieniecka-Gazarkiewicz et al., 2021) was strongly upregulated during recovery. PtLPCAT1 has been recently shown to be localized to the chloroplast ER membrane (Huang et al. 2023) and its activity to affect both TAG and galactolipid synthesis (You et al., 2022). The expression of two genes encoding cytoplasmic long-chain acyl-CoA synthetases *ACS4* and A*CS2* (Phatr3_J45510 and Phatr3_J12420, respectively) (Jallet et al., 2020), catalyzing the activation of free fatty acids by CoA, was upregulated after N resupply. Upregulated expression of these genes is consistent with their role in delivering acyl-CoA for the extraplastidial membrane lipid synthesis and possibly for the degradation *via* β-oxidation. Lipid traffickers, such as phospholipid flippases and scramblases, are enzymes that move phospholipids from one leaflet of the membrane to another (López-Marqués, 2021). Interestingly, high mRNA levels of phospholipid flippase (Phatr3_J52368) and phospholipid scramblase (Phatr3_J55111) genes were found under both N starvation and recovery.

**Table 1.** Upregulated lipid metabolism-related genes (other than lipases) 24 h after N resupply (recovery)

| Classification | Gene ID | adjusted p value | Localization | References |
| --- | --- | --- | --- | --- |
| Galactolipid synthase | Phatr3_J54168/MGD1 | 4.79E-142 | Plastid | Shang et al., 2022 |
|  | Phatr3_J43116/ DGDG1/2 | 7.39E-15 | Plastid |  |
| Sulfolipid synthase | Phatr3_J21201/SQD1 | 2.11E-64 | Plastid |  |
| Acyltransferase | Phatr3_J20460/LPCAT | 1.50E-167 | PPC*, ER | Jasieniecka-Gazarkiewicz et al., 2021; Huang et al. 2023 |
| Phospholipid flippase | Phatr3_J52368/ALA | 5.84E-26 | Membrane |  |
| Phospholipid scramblase | Phatr3_J55111/PLS1 | 4.10E-185 | Membrane |  |
| Acyl carrier protein | Phatr3_J9709/ | 4.19E-78 | Plastid |  |
| Acyl-CoA synthetases | Phatr3_J45510/ ACS4 | 1.26E-165 | Cytoplasm | Jallet et al., 2020 |
|  | Phatr3_J12420/ ACSL | 3.47E-36 | Cytoplasm | Jallet et al., 2020 |

\*PPC - the periplastidial compartment

## Discussion

Nitrogen starvation stress in *P. tricornutum* is manifested by the extensive dismantling and degradation of organelles, including plastids, concomitant with the formation and sequestration of TAG into the LDs. Upon the cessation of N starvation by N resupply, LDs are broken down and their TAG core is degraded by lipases and possibly by lipophagy — the autophagic degradation of LDs. ABPP has successfully validated the role of serine hydrolases in lipid metabolism across animal, plant, and bacterial systems (Ruby et al., 2017; Bassett et al., 2018; Dolui and Vijayaraj, 2020). However, the SH family in diatoms, and generally in microalgae, is largely uncharacterized and unexplored by the functional chemoproteomics approach. Therefore, a comprehensive landscape of the *bona fide* active SHs was elucidated using a gel-based competitive ABPP and a gel-free ABPP approaches. The activity profile of most of the identified mSHs and serine proteases generally correlated with the transcript level of the genes encoding these proteins. Importantly, none of these identified SHs have previously been characterized, to our knowledge, underscoring the potential of ABPP as the tool of choice in determining the lipid catabolic linchpins. Many of these lipases were enriched in the sample processed from the “recovery” cells. These functionally relevant enzymes can be prioritized for further functional characterization.

### Phenyl mercuric acetate: an SH (lipase) inhibitor

The competitive ABPP has a proven record in developing inhibitors for various mSHs (Leung et al., 2003) with exceptional specificity and selectivity, since inhibitors are tested against many enzymes in parallel. We report PMA as a potential SH inhibitor that was experimentally validated in the competitive ABPP assay. PMA was previously reported to suppress fungal lipase activity (Sharma, 1976), wheat galactolipase (O’Sullivan et al., 1987), glucocerebrosidase, and α-galactosidase-hydrolases involved in glycolipid metabolism (Motabar et al., 2010). SHs often exhibit moonlighting activity; for example, a fungal cutinase was reported to possess both galactosidase and lipase activities (Verma et al., 2013). We speculate that the SH inhibited by PMA in the competitive ABPP might be a putative multifunctional lipase that can catalyze lipolysis of a broad range of lipids. Lipidome profiling confirmed that N-starved cells treated with PMA during recovery were unable to mobilize TAGs, degrade LDs and restore photosynthetic apparatus. A similar pattern of changes in the molecular species of DGDG, PC, and DAG (**Fig. 3**) was noted, manifested by a higher abundance of 36:6 (20:5/16:1) species, was found in the control untreated sample. We speculate that the lipid remodeling route PC→DAG→DGDG during recovery, likely involving the LPCAT and phospholipases, may be inhibited in the PMA-treated cells. Therefore, the cells were unable to remobilize storage lipids and reestablish a functional plastid. It should be noted that *P. tricornutum* did not survive the high PMA concentrations used in the *in vitro* assay (**Fig. S2**), possibly due to the inhibition of photosynthetic and other primary activities at a higher concentration of the inhibitor (Honeycutt and Krogmann, 1972).

### ABPP unlocked differential activomes in *P. tricornutum*

The phenomena of storage lipid accumulation and mobilization are well balanced in *P. tricornutum* to ensure cellular energy homeostasis. This is assisted by the differential activity of SHs, which facilitate optimized nutrient (C and N) and energy expenditure. In contrast to N starvation, when active SHs were enriched in hydrolases likely involved in the degradation of membrane lipids and chlorophyll, recovery from N-starvation was marked by a different set of mSHs dominated by lipid hydrolases. Nitrogen stress in diatoms and microalgae is characterized by the induction of proteases, attributed to scavenging and remobilization of N from proteins for the synthesis of proteins essential for survival during N deprivation (Berges and Falkowski, 1998). For example, the Deg-P protease (B7G9W1), active under N starvation, is predicted to have serine-type endopeptidase activity (Family S1) and mediate proteolytic cleavage of the chloroplast proteins. The Deg-P protease is involved in cleavage of the D1 protein of photosystem II (Kanervo et al., 2003) and in the elimination of misfolded proteins. This protease seems to play a pivotal role in *P. tricornutum* during massive plastid ablation under N starvation. N-starvation is also marked by the loss of chlorophyll (chlorosis). Chlorophyllase is the first enzyme in the chlorophyll degradation pathway (Shioi et al., 1991) and may also catalyze the transesterification of chlorophyllide. We identified two functional chlorophyllases active during N starvation. Two subtilin proteases (B7G7C5, B7FXC9) were exclusively enriched during recovery. Subtilases are ubiquitous plant proteases, playing also a pivotal role as processing proteases (Schaller et al., 2018). The role played by these subtilases in Phaeodactylum is elusive. The two most abundant SHs found in this study were S10 subfamily serine carboxypeptidases with a highly conserved α/β hydrolase fold. Similarly, in a benthic diatom *T. pseudonana*, both the protein, and the transcript abundance of the serine carboxypeptidase gene were increased twofold under N starvation (Hockin et al., 2012).

### Lipases: the linchpins in lipid catabolism

Metabolic SHs cleave the ester, amide, or thioester bonds in small molecules, peptides, or proteins using a conserved serine nucleophile. Lipases are the major catalytic members of the mSH family. We found a unique enrichment of SHs that are likely involved in processing of a broad spectrum of lipids. This lipid hydrolase oligarchy prevailed during TAG remobilization. A functional phospholipase/carboxylesterase/thioesterase domain-containing protein (B7G4G0), despite its annotation as lysophospholipase, shows structural similarity to acyl protein thioesterase APT 2. APT are members of the SH family and can be detected by ABPP. APT 2 was shown to play role in disintegration of the lipid bilayer by removing acyl chains from its protein substrate (Abrami et al., 2021).

Once accessed by TAG lipase, the lipid core is rapidly depleted to ensure the supply of acyl groups for membrane lipid synthesis and for energy and glycogenesis through beta-oxidation. Two such potential class 3 lipases (B7FZK6, B7FV70) were found to be active exclusively during recovery. B7FV70 shows similarity to lysosomal acid lipase, which is involved in the hydrolysis of TAG and cholesteryl esters (Li and Zhang, 2019). One of the major mSHs (B7G4E7/PHATRDRAFT_47650) identified in this study is annotated as a “Lipase_3 domain-containing protein. It contains the AB_hydrolase domain, bears the consensus lipase sequence GxSxG, and is predicted to play a role in lipid metabolism by Gene Ontology (GO) enrichment analysis. One can extrapolate from this finding that ABPP should be the method of choice for investigating uncharacterized SHs, regardless of their degree of annotation (Simon and Cravatt, 2010). These findings also include a carbohydrate esterase (B7G104), which belongs to the SGNH superfamily. This superfamily’s enzymes exhibit broad substrate specificity and catalyze diverse substrates ranging from glycolipids to glycoproteins. This is a notable finding since *P. tricornutum* harbors glycolipids, such as SQDG, and betaine lipids, including DGTA, in the monolayer of lipid droplets (Guéguen et al., 2021). An S-formyl glutathione hydrolase (B7FP91) belonging to esterase D of the SH superfamily was found exclusively during recovery. This enzyme exhibits carboxyl esterase activity and may be involved in the hydrolysis of S-formylglutathione to glutathione and formate (Haslam et al., 2002). As mentioned before, certain lipases, not detected by ABPP, may represent enzymes that have highly restricted cellular distribution or ability to recognize and react with the probe (Bachovchin et al., 2010).

The enrichment of a few low-abundance lipases was revealed exclusively by activity-based tagging, emphasizing that high throughput functional proteomics, such as active site peptide profiling, is sufficiently sensitive in detecting functionally relevant proteins irrespective of their relative abundance. Unlike abundance-based proteomics, enrichment of probe-labeled enzymes by streptavidin agarose prior to LC/MS analysis greatly increases the probability of identifying target proteins (Okerberg et al., 2005; Dolui and Vijayaraj, 2020). For example, a lipase (B7FV70/Phatr3_J44989) and phospholipase (B7FTN8**/**Phatr3_J33720) have low levels of mRNA expression in our transcriptomics data. However, the active proteins were successfully detected by activity-based tagging. Further, there was a poor correlation between mRNA and the chlorophyllase protein (Phatr3_J43025) as it was downregulated under N starvation but was enriched by activity-based peptide profiling, pointing to a major discrepancy between the mRNA-protein correlations. Many variables such as mRNA decay, translation, and protein degradation rates often lead to this poor correlation (Ponnala et al., 2014).

### Dynamic metabolic SHs (mSHs) in N- starvation

The chlorophyllase found is an atypical member of the SH superfamily, which corroborates the expanding diversity of the SH activity repertoire. Indeed, chlorophyllase, a serine hydrolase, was previously reported in *Chenopodium album* (Tsuchiya et al., 2003). We also revealed an abundant amidase (B7GE02), which has been previously found in the proteomics study of the N-limited *P. tricornutum* (Burch et al., 2021). Another notable finding from the gel-free ABPP was the enrichment of an AB hydrolase superfamily protein B7GCV4. According to the existing annotation, it belongs to the MetX_acyltransf family and is predicted to have transferase activity along with its prototypical hydrolase activity. Recently, human ABHD 14B was shown to be an atypical SH, functionally annotated by ABPP as a lysine deacetylase, which transfers an acetyl group to coenzyme A (Rajendran et al., 2019). These mSHs were functional exclusively under N starvation, which was also corroborated by the transcriptomics data, pointing to the critical and indispensable roles played by these enzymes under these conditions.

### Phospholipases: The nemesis of membrane lipids under N starvation

Transcriptomics data provided a complementary insight into the role of putative phospholipase classes in lipid remodeling and providing precursors for TAG biosynthesis. Based on their expression profile, patatin-like phospholipases and phosphodiesterases (phospholipase C & D) seem to be the main driving force behind membrane lipid dismantling in *P. tricornutum* during N starvation. Patatin-like lipases may exhibit diverse activities and cleave acyl groups from triacylglycerols, membrane lipids, including phospholipids (Mignery et al., 1988) and galactolipids (Li et al., 2013). We suggest that some patatin-like PLPs showing an increased gene expression under N starvation, may be involved in the release of acyls groups from phospholipids and the generation of glycerol-3-phosphodiesters. Consecutive processing by phospholipases C and D liberates the precursors for TAG biosynthesis. However, the role of phospholipase D may extend beyond phospholipid degradation. Recently, Arabidopsis phospholipase D has been reported to be involved in autophagy in response to N deficiency by hydrolyzing the ATG8-PE conjugate (Yao et al., 2022). We hypothesize that phospholipase D may play a similar role in autophagy in *P. tricornutum.* Interestingly, the gene encoding the hallmark autophagy protein ATG8 (Phatr3_J50303) was also found to be significantly upregulated under N starvation in our transcriptome. It has a signature phosphatidylethanolamine-amidated glycine for lipidation. Therefore, we speculate that it may play a role in yet-to-be studied autophagy in *P. tricornutum*.

An opposite trend of expression was found for genes encoding enzymes involved in membrane lipid synthesis and restoration of the plastid membranes (Table 1). The genes encoding lipid traffickers, such as flippases and scramblases, which play role in redistributing phospholipids in various cell membranes were also strongly upregulated.

### Acyl hydrolases and lipid homeostasis

Sixteen class 3 lipases were enumerated from the genome mining of *P. tricornutum* databases. The majority of these class 3 lipases did not show the conspicuous changes in gene expression under our experimental conditions except for four genes (Phatr3_J43856, Phatr3_J44028, Phatr3_J45895, Phatr3_J50397), which were significantly upregulated during recovery from N-starvation (**Fig. 6**) and therefore may play role in TAG degradation. Notably, two genes (Phatr3_J43856, Phatr3_J44028) were highly expressed under N-starvation, suggesting their involvement in TAG turnover for energy and carbon homeostasis under this condition. In line with this premise, Phatr3_J43856, is annotated as a gene for the autophagy-related lipase ATG15, involved in vacuolar breakdown of autophagic body membrane and TAG breakdown (lipophagy) in yeast (Hirata et al., 2021). Further study on the role of this lipase in *P. tricornutum* is needed. The significant upregulation of Phatr3_J44028 under N and P deprivation was also reported in many studies (Cruz de Carvalho et al., 2016; Matthijs et al., 2015; Matthijs et al., 2017) placing this lipase as the principal lipase involved in TAG homeostasis. The gene for an activating protein of TAG lipase CGI58 (Phatr3_J54974), which possesses the AB hydrolase domain, was downregulated during recovery but strongly expressed in N starvation, supporting its role in TAG homeostasis during N starvation. A recent study has shown that PtCGI-58 plays a role in the recycling of acyl chains derived from mitochondrial β-oxidation into TAG biosynthesis (Shu et al. 2023).

### Conclusions

This study provides a new perspective on lipid metabolism in *P. tricornutum*, highlighted by the identification and differential activity of SHs. This work revealed a much-needed profile of active SHs in this diatom, emphasizing lipases as major metabolic serine hydrolases involved in the remobilization of storage lipids during recovery from N starvation. Overall, the study provides a better template for comprehending lipid metabolism in *P. tricornutum* and provides further stimuli for basic and translational research. The approach described here can also be applied to other microalgae and diatoms.

## Materials and Methods

### Chemicals and reagents

ActiveX TAMRA-FP (cat no. 88318) and ActiveX Desthiobiotin-FP (cat no. 88317), Zeba spin desalting columns, 5 ml (cat no. 89892) were procured from Thermo Scientific (USA). Streptavidin agarose (cat no. S1638), dimethyl sulfoxide (cat no. W387520), dithiothreitol (cat no. D9779), iodoacetamide (cat no. I6125), urea (cat no. U5378), and MS grade trypsin (cat no. T6567) were purchased from Sigma-Aldrich (USA).

### Phaeodactylum tricornutum cultivation

*Phaeodactylum tricornutum* strain UTEX 646 (Pt4) was cultivated in a Red Sea Salt Enriched (RSSE) medium (Taparia et al., 2022). The 100-mL cultures were maintained in 250-mL Erlenmeyer flasks in an incubator shaker at 22 °C and 150 rpm with a light intensity of 50 μmol photons m^−1^·s^−1^ and a CO2-enriched atmosphere of 100 mL CO2·min^−1.^ Weekly dilution was performed to maintain the nutrient-replete growth. Cultures were subjected to N starvation by excluding KNO3 from the RSSE medium. For recovery from N starvation, cells were harvested and resuspended in an N-replete RSSE medium (designated as a recovery medium).

### Protein extraction

Cells from 100-mL cultures were collected after 120 h of N starvation conditions and 24 h of recovery by centrifugation at 3000 rpm and washed once with PBS. The cell pellets were resuspended in 1.5 mL lysis buffer [50 mM Tris-HCl (pH 8.0), 150 mM NaCl, 1mM MgCl2, 1mM KCl, 10% Glycerol] and vortexed for 30 min at 4 °C in the presence of glass beads (0.5- mm diameter). The cell debris was removed by centrifugation at 6,000 rpm for 10 min at 4 °C. Total protein concentration estimation in cell-free lysates was carried out by the Bradford method (Bradford, 1976).

### Gel-based ABPP and in vitro competitive ABPP

Labeling of serine hydrolases was performed with 2 μM of FP-serine hydrolase probe. In brief, 40 µg of protein was pre-incubated with either 100 μM of inhibitor (PMSF, Fenitrothion and Phenyl Mercuric Acetate) DMSO for 30 min at 37 °C. Next, a 2-μM probe was added into each sample, and incubation was extended for another 1 h at 37 °C in the dark. The reaction was terminated by the addition of 4x loading buffer (200 mM Tris-HCl, pH 6.8, 400 mM DTT, 8% SDS, 0.04% bromophenol blue, and 40% glycerol) and boiled at 95 °C for 5 min. Proteins were resolved on 12% (w/v) SDS-PAGE, and the labeled enzymes were detected using Cy3 /TAMRA settings (excitation wavelength 532 nm) in a Typhoon FLA9500 phosphor imager (GE Healthcare Life Sciences). Further, the gels were stained with Coomassie Brilliant Blue R-250, and the images were documented.

### *In vitro* competitive ABPP

A stock solution (0.5 mM) of Phenyl Mercuric Acetate (PMA) was freshly prepared in DMSO. For concentration-dependent inhibition, three concentrations of PMA were used. A control was also included wherein only DMSO was added. To check *bona fide* inhibition of lipase activity, the competitive ABPP with *P. tricornutum* cells 24h after recovery from N-starvation was performed. In brief, PMA (25, 50, and 100 nM) was added to 20 mL- cultures at an initial cell density of 2.0 x 10^6^ cells/mL. Growth of *P. tricornutum* cultures was monitored by cell counting using a Luna cell counter (BioCat, GmbH) at every 24-h interval. Lipid droplets were observed using a Zeiss Axio Imager A2 fluorescence microscope (Carl Zeiss MicroImaging Inc., Göttingen, Germany), and images were photographed. For this, 1 µL of Nile Red (0.1 mg/mL), prepared in DMSO, was added to a 10-µL cell suspension.

### Enrichment of serine hydrolase followed by on-bead trypsin digestion

This experiment was performed in three biological replicates. In brief, one microgram (500 µL) of protein was incubated with 10 µL (20 µM) of a desthibiotin conjugated FP-Probe (COMPANY). A control was also included whereby the probe was replaced with 10 µL of DMSO. Labeling was performed at 37 °C for 1 h. The reaction was stopped by adding 500 μL of 10 M of urea, followed by reduction with 10µL of 500 mM DTT (incubation at 65 °C for 30 min) and alkylation with 40 µL of 1M iodoacetamide in the dark. Desalting was performed using a Zeba Spin desalting column (Thermo Scientific). Labeled proteins were enriched with 100 μL of streptavidin agarose for 1 h at room temperature in an end-to-end tube rotator. Subsequently, the beads were separated by centrifugation at 1000 g for 1 min. The unbound proteins were removed by washing the beads thrice, each time with 1.5 mL of washing buffer [25 mM Tris, (pH 8.0), 300 mM NaCl, 1% NP 40], PBS, and LC-MS grade water, respectively. The affinity captured proteins were then digested with 4 μg of MS grade trypsin (Sigma-Aldrich) in 500 μL of digestion buffer [2 M urea, 20 mM Tris, (pH 8.0)] for 16 h at 37 °C with constant shaking. The digested peptides were collected by centrifugation at 1000 g for 1 min, and the supernatant was dried in a vacuum concentrator (Eppendorf). The tryptic peptides were acidified to 0.1% formic and desalted using C18 tips, dried and re-suspended in 0.1% formic acid.

### Active site peptide profiling

Active site peptide fingerprinting of serine hydrolases was carried out as described by (Dolui and Vijayaraj, 2020).

### LS/MS proteome analysis

The peptides were resolved by reverse-phase chromatography on 0.075 × 300-mm fused silica capillaries (J&W) packed with Reprosil reversed phase material (C18-AQ 3µm, Dr Maisch GmbH, Germany). The peptides were eluted with a linear 60-min gradient of 5 to 28%, followed by a 15-min gradient of 28 to 95% and 25 min at 95% acetonitrile with 0.1% formic acid in water at a flow rate of 0.15 μL/min. Mass spectrometry was performed by a Q Exactive HF mass spectrometer (Thermo) in a positive mode using a repetitively full MS scan followed by collision-induced dissociation (HCD) of the 18 most dominant ions selected from the first MS scan. Proteomics was performed at the Smoler Proteomics Center (Technion, Israel).|

### Data analysis

The mass spectrometry data were analyzed using Protein Discoverer 2.4 (Thermo) using the Sequest search engine, searching against the *Phaeodactylum tricornutum* database, and a decoy database. Peptide- and protein-level false discovery rates (FDRs) were filtered to 1% using the target-decoy strategy. Oxidation on methionine, protein N-terminus acetylation, and biotin on serine (Thermo-88317) were accepted as variable modifications, and carbamidomethyl on cysteine was accepted as static modifications.

### RNA sequencing and data analysis

N-starved cells (120h-N) and cells after 24h of recovery in replete medium (24h+N) were processed for RNA isolation and transcriptomic analysis. Total RNA was extracted using an SV Total RNA Isolation System Kit (Promega). The integrity of isolated RNA was evaluated on an Agilent 2100 Bioanalyzer. Messenger RNA (mRNA) libraries were prepared for each sample and sequenced using the Illumina HiSeq-2000 platform. The RNAseq reads were quality and adapter trimmed with Trim_galore v0.6.7 (Cutadapt v3.7 and FASTQC v0.11.9) [,github.com/FelixKrueger/TrimGalore], mapped to the *Phaeodactylum tricornutum* genome v2 (ASM15095v2, Ensembl Protists release 53; protists.ensembl.org/Phaeodactylum_tricornutum/Info/Index) using STAR v2.7.10a (Dobin et al., 2013) and differential expression was evaluated using DESeq2 v1.36.0 (Love et al., 2014) (absolute log2 fold change ≥ 1 and adjusted *p* value < 0.05). Functional annotation of the genome was performed using Trinotate v3.2.1 [github.com/Trinotate/Trinotate.github.io/wiki]. Then the functional enrichment of differentially expressed genes was performed using topGO v2.48.0 (Alexa and Maintainer, 2021) topGO: Enrichment Analysis for Gene Ontology. R package version 2.48.0] (Fisher’s exact test).

### Fatty acid methyl ester (FAME) analysis

Fatty acid methyl ester analysis of lipids was performed as described by (Taparia et al., 2022). In brief, 50 x 10^6^ cells were collected every 12 h during recovery from N stress and freeze dried. Freeze-dried samples were transmethylated with 2% (v/v) H2SO4 in anhydrous methanol at 80°C for 1.5 h under an argon atmosphere with continuous stirring. Pentadecanoic acid (C15:0) (Sigma-Aldrich) was included as an internal standard. FAMEs were quantified on a Trace GC Ultra (Thermo, Italy) equipped with an FID.

### Lipid extraction

Total lipids were extracted from N-replenished cells (24 h in the nitrogen recovery medium). Briefly, 150 million cells from five biological replicates of the control and five biological replicates of PMA (100 nM) treated cells were collected and processed for lipid extraction. Cells were subjected to hot isopropanol pre-heated at 70 °C for 10 min. Lipid extraction was carried out with methanol: chloroform (2:1) with constant mixing under an argon atmosphere for 1 h at RT. The sample was centrifuged, and the solvent was collected. This step was repeated, and all solvent samples (including isopropanol) were pooled together in a single tube.

Double distilled water was added and mixed by vortexing, and phase separation was achieved by centrifugation. Finally, the organic phase was collected and dried under an N2 stream flow.

### Untargeted lipidome profiling by Liquid Chromatography-Mass Spectrometry

Prior to LC-MS analysis, samples were re-suspended in 70% (v/v) acetonitrile and 30% (v/v) isopropanol. LC–MS data were obtained using a Waters Acquity UPLC system equipped with a C8 reverse-phase column (Waters), coupled to a q-Exactive mass spectrometer (Thermo Fisher). To acquire mass spectra, a mass range from 100 to 1500 m/z was included. Resolution was set to 70,000, with three scans per second. Capillary voltage was set to 3 kV, with a sheath gas flow value of 60 and an auxiliary gas flow of 20 (values are in arbitrary units). Capillary temperature was set to 275 °C; and the temperature of the drying gas in the heated electrospray source to 300 °C. Chromatograms from the UPLC–FT/MS runs were analyzed and processed with REFINER MS 11.0 (GeneData). The following information was obtained from each chromatogram: retention time, molecular masses, and associated peak intensities. The chemical noise was subtracted automatically. To align the chromatograms, we used a pairwise alignment-based tree approach. This involved defining m/z windows of five points and retention-time windows of five scans within a sliding frame of 200 scans. The further processing of the MS data included isotope clustering, adduct detection, and a library search. The method we used could not identify DGTA, a major betaine lipid in *P. tricornutum*. Resulting data matrices with peak ID, retention time, and peak intensities in each sample were generated. Batch-normalization and sample-median-normalization were conducted, and the resulting data matrices were used for further analysis. Data were processed at the Metaboanalyst platform (https://www.metaboanalyst.ca/) for statistical analysis.

## Supporting information

SRA metadata

Supplemental File S1 Recovery and starvation Active Site Peptide Profiling

Supplemental File S2 Medium_recovery_vs_Nitrogen_deseq_results_genes_padj

Supplemental file S3 GO terms

ALL Supplemental Figures

## Acknowledgements

This research was supported by the grant from Israeli Science Foundation (grant No1992/21)

