## Supplementary material for "Chemoproteomic profiling of serine hydrolases reveals the dynamic role of lipases in *Phaeodactylum tricornutum*": ALL Supplemental Figures

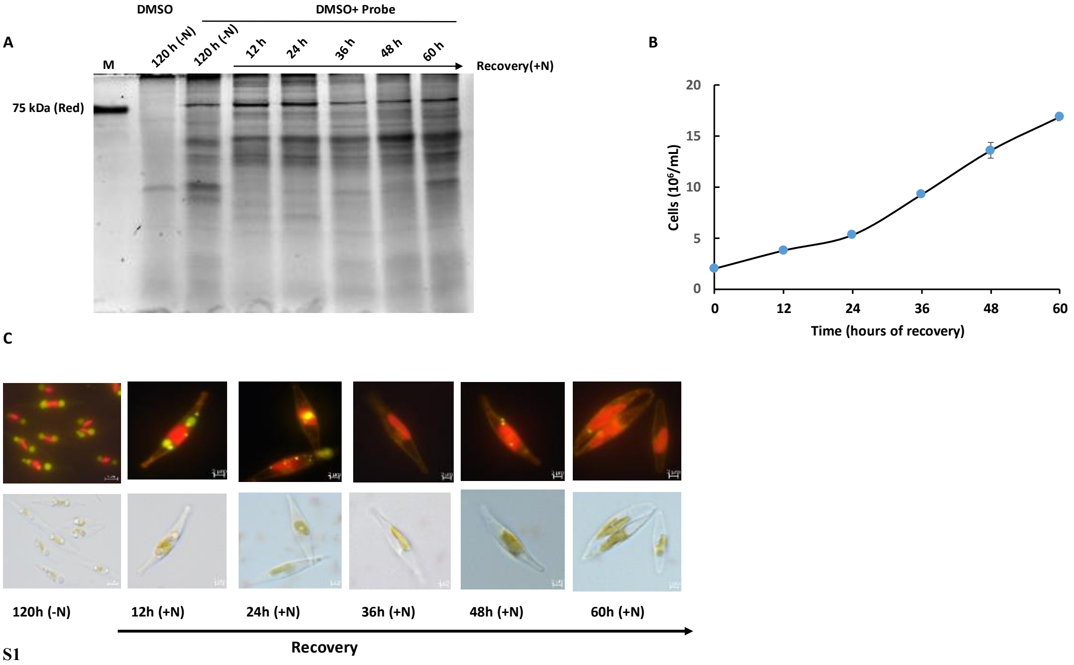


Fig. S1

**Figure S1. SH activity detected by in gel-ABPP during lipid droplet accumulation and remobilization.** *P. tricornutum* cells were monitored during lipid droplet accumulation under N starvation and during remobilization of lipid droplets upon nutrient replenishment (Recovery). (**A**) In-gel fluorescence ABPP of the proteome processed from N-starved cells (120h-N) and during recovery every 12 hours. (B) Cell numbers during 60h of recovery from N starvation. (C) Micrographs of *P. tricornutum* cells after 120 h in N starvation and during recovery. Upper raw: culture aliquots (10 µL) were stained with 1 µL Nile Red solution (0.1 mg/mL in DMSO) to visualize lipid droplets using a fluorescence mode; Bottom raw: cells observed using a Differential Interference Contrast (DIC) mode.

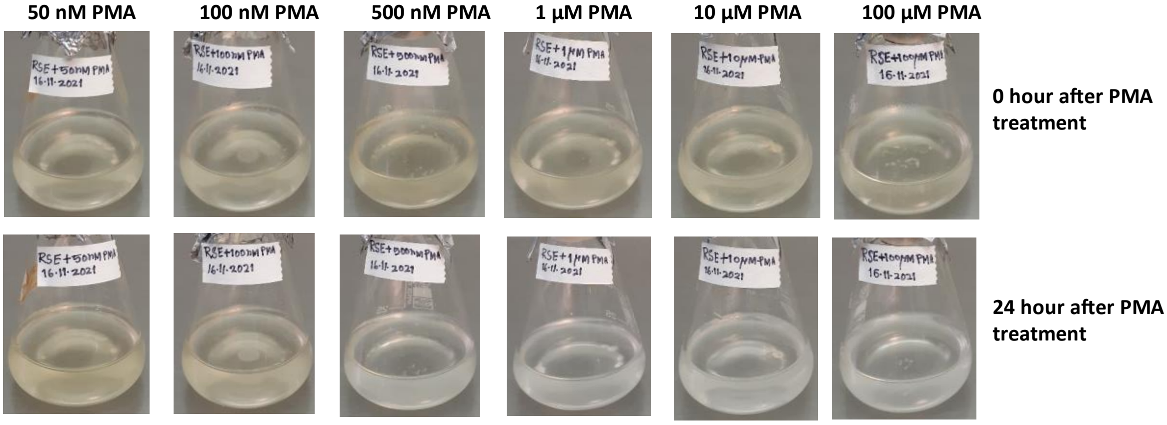


**Figure S2. Concentration-dependent inhibition of *P. tricornutum* cells during recovery from N starvation.** Six concentrations of PMA (from 50 nM to 100 µM ) were tested to evaluate the sensitivity *P. tricornutum* cells and the degree of *in vivo* inhibition on lipolytic activities as manifested by visible growth. Cultures with the initial cell density of 2 106 cell/mL were treated with PMA. Upper panel: images of *P. tricornutum* flask cultures at 0 h of PMA treatment. Lower panel: after 24 hour of recovery, growth is visible only with 50 and 100 nM PMA.

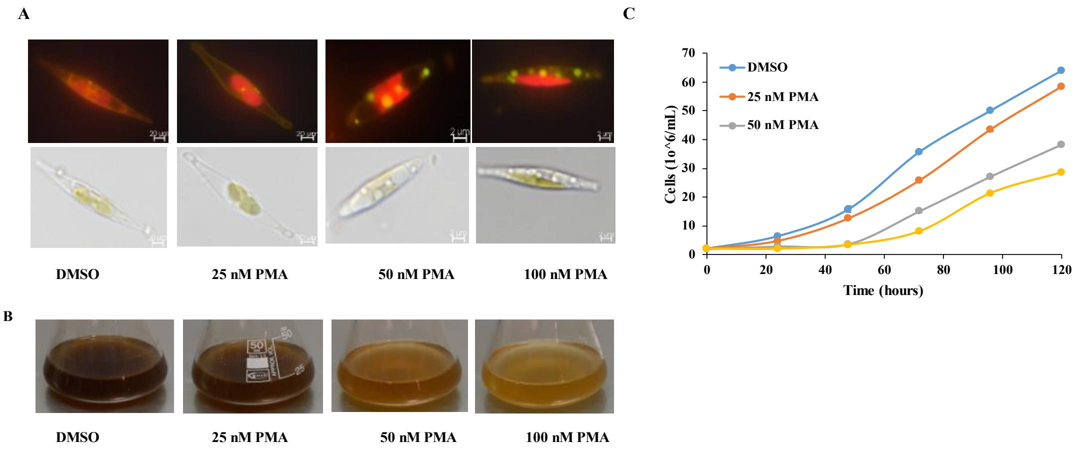


**Figure S3.** ***In vivo*** **effect of PMA in** ***P. tricornutum*** **cells in N replete medium.**

Cultures inoculated at the cell density of 2 x $10^6$/mL, were treated with three concentrations of PMA (25, 50 and 100 nM) and growth was monitored during 120h. Control (DMSO-treated cells) as PMA was dissolved in DMSO. Growth performance and degradation of lipid droplets in the presence of PMA were examined every 24 h. (**A**) Micrographs of *P. tricornutum* cells with lipid droplets stained with Nile Red. Images were taken from control and PMA-treated cells after 48h. (**B**) *P. tricornutum* cultures after 72 h. (**C**) Growth curves of control and PMA-treated cultures.

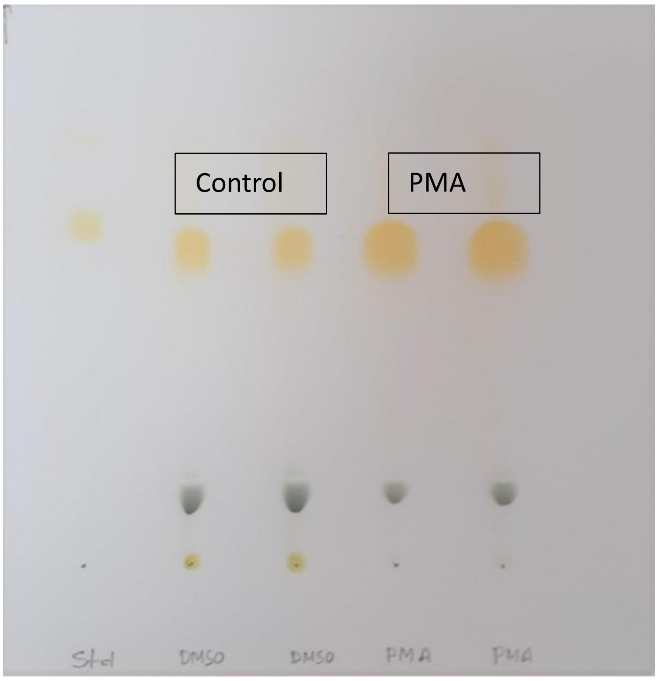


**Figure S4.** TLC separation of lipid extracts of control and PMA treated cells used for lipidomics analysis in a solvent system for neutral lipids. TAG spots are visible after iodine staining.

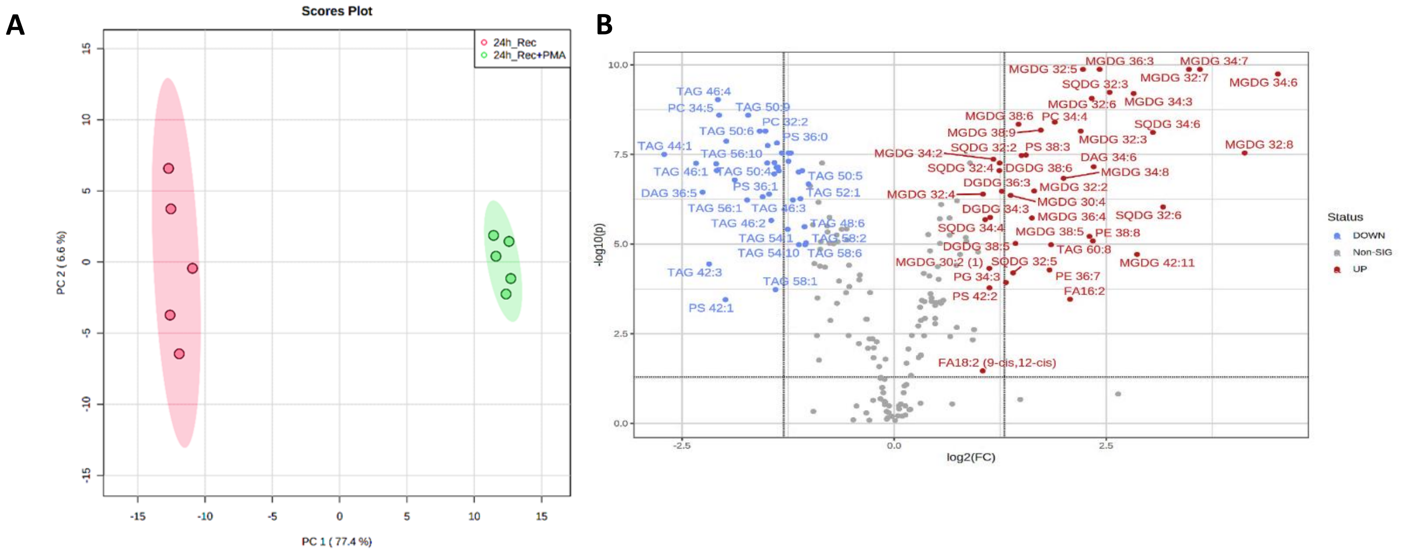


**Figure S5.** Effect of PMA treatment on the lipidome of *P. tricornutum* during recovery from N starvation. (**A**) A principal component analysis (PCA); (**B**) Volcani plot represents the distribution of significantly different lipid species in PMA-treated cells: significantly more abundant lipids highlighted in red, significantly less abundant species highlighted in blue.

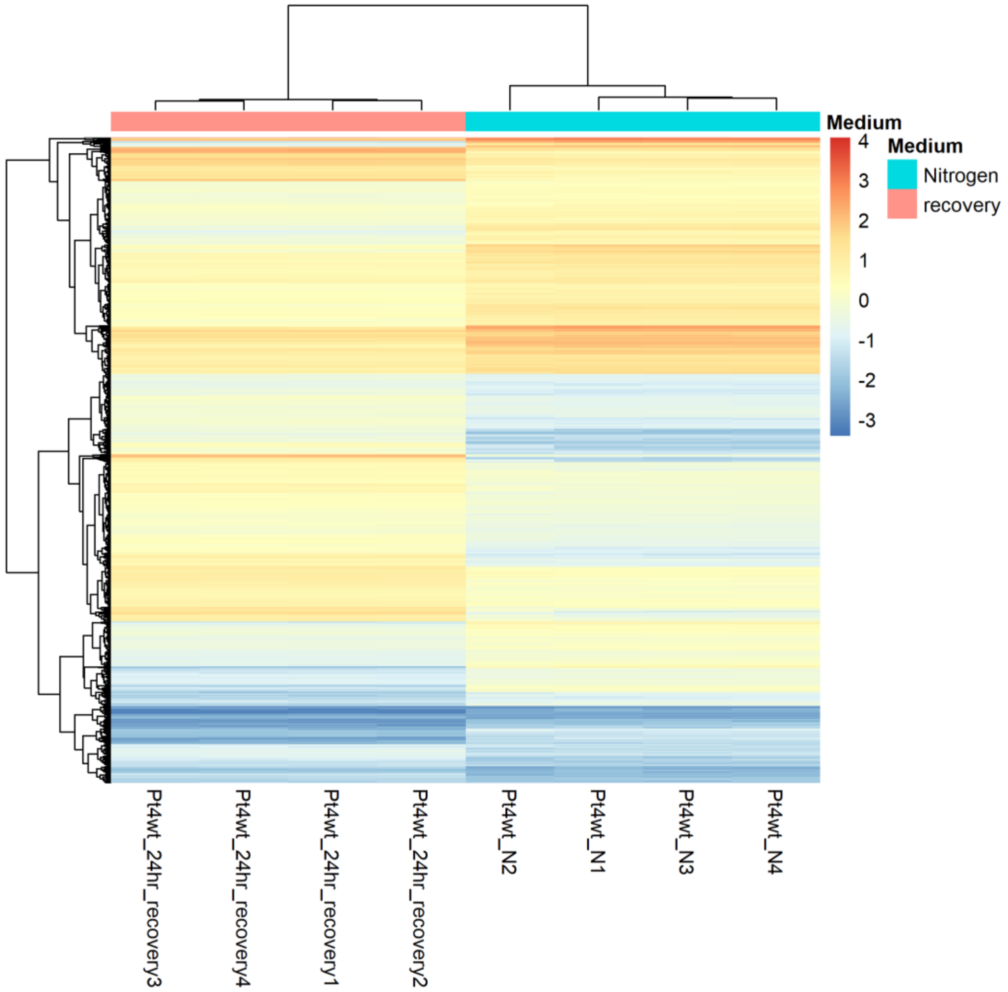


**Figure S6.** A heatmap represents differential expression levels of mRNA between 120h of N starvation and 24h of recovery. Module color is displayed.
